# BAV-LLPS: A database of bacterial, archaea and virus liquid-liquid phase separation proteins

**DOI:** 10.1101/2025.08.15.670539

**Authors:** Cecilia B. Rodriguez, Ronaldo Romario Tunque Cahui, Nicolas Demitroff, Layla Hirsh, Damien P. Devos, Graciela Boccaccio, Gustavo Parisi

## Abstract

Liquid–liquid phase separation (LLPS) is a key process underlying the formation of biomolecular condensates (BMCs), such as membrane-less organelles (MLOs), that compartmentalize biochemical processes inside the cells. While LLPS has been extensively studied in eukaryotes, its role in bacteria, archaea, and viruses remains far less characterized. Recent studies in bacteria have revealed that LLPS-driven condensates play critical roles in RNA processing, stress response, and pathogenicity. Similarly, many viruses exploit LLPS to facilitate crucial steps in their infection cycles, including viral entry, genome replication, assembly, and host immune evasion.

In this work, we introduce a hand-curated database of LLPS proteins from bacteria, archaea, and viruses (BAV-LLPS Database). This resource, extended through sequence similarity searches, comprises over 5,000 proteins and integrates diverse data including biological annotations, sequence features, predicted disordered regions, LLPS per site probability, and AlphaFold2-based structural models. Additionally, our web server enables users to explore both the curated and homologous derived datasets, providing a platform to uncover evolutionary relationships and intrinsic and differential properties of LLPS proteins across various taxonomic groups. This work seeks to deepen our understanding of LLPS mechanisms beyond eukaryotic organisms, emphasizing their significance across diverse life forms. It also aims to foster the development of specialized predictive tools that will facilitate the exploration and characterization of LLPS processes in a wide array of living organisms, thereby contributing to advancements in both fundamental biological research and applied biomedical sciences.

**Availability and Implementation:** BAV-LLPS DB is freely accessible at

https://bav-llps-db.bioinformatica.org/. The data can be retrieved from the website. The source code of the database can be downloaded from https://bav-llps-db.bioinformatica.org/download

## Introduction

Liquid-Liquid phase separation (LLPS) involves the transition from an homogeneous system containing given macromolecular components into two separated phases, one enriched and another depleted in the involved macromolecules (Alberti and Dormann, 2019). LLPS is greatly influenced by intrinsically disordered protein regions (IDRs) (Fonin *et al*., 2022; Tesei *et al*., 2021) and generates the so-called biomolecular condensates (BMCs) or membrane-less organelles (MLOs) (Fernández-Alvarez *et al*., 2023; Darling and Uversky, 2023). Unlike other eukaryotic organelles such as mitochondria or Golgi apparatus, BMCs and MLOs lack a delimiting lipid membrane. Due to their reversible and dynamic formation, BMCs and MLOs are transient microcompartments that can selectively concentrate certain macromolecules while excluding others. In doing so, MLOs generate microenvironments that can accelerate or inhibit biochemical reactions, they can buffer protein and nucleic acid concentrations, participate in the sensing of environmental changes, control signaling pathways, and in the nucleation of cytoskeletal structures among other functions. Dozens of BMCs and MLOs have been identified in mammalian cells, a list which is continually expanding(Fernández-Alvarez *et al*., 2023; Darling and Uversky, 2023; Thomas *et al*., 2025). Proteins undergoing LLPS and their derived condensates have been found in different organisms across eukaryotes, prokaryotes, and viruses reflecting an evolutionary conservation of these mechanisms to control metabolic processes using spatial and temporal compartmentalization (Rangachari, 2023). In addition to their broad taxonomic distribution, LLPS-related proteins in bacteria, archaea, and viruses are significantly less characterized than their eukaryotic counterparts. In fact, established databases dedicated to LLPS proteins (i.e PhaSepDB (You *et al*., 2020), LLPSDB (Li *et al*., 2020) and MLOsMetaDB (Orti *et al*., 2024)) are predominantly populated with eukaryotic examples.

In contrast to the well characterized eukaryotic MLOs and BMCs, interest in prokaryotic BMCs has risen abruptly in recent years. The use of *in vitro* reconstitution, fluoroscopic assays, differential interference contrast microscopy and super-resolution microscopy applied to *in vivo* studies, have significantly advanced the detection of proteins associated with LLPS in bacteria, facilitating the exploration of the importance of BMCs in prokaryotes (Azaldegui *et al*., 2021). One of the systems better characterized in prokaryotic condensates are the bacterial ribonucleoprotein bodies (BR-bodies), which control mRNA decay by locally concentrating the RNA degradosome proteins and their substrates (Al-Husini *et al*., 2018); (Al-Husini *et al*., 2020). Different proteins participating in LLPS in bacteria were also related to different biological processes involved with the pathogenicity, resistance to antibiotics or antimicrobial agents and bacterial survival (Liu *et al*., 2024). In the human commensal *Bacteroides thetaiotaomicron*, the activity of the transcription termination factor Rho largely depends on its LLPS, with consequences in gene expression and bacteroid competence (Krypotou *et al*., 2023). Another well characterized bacterial condensate involves the proteins FtsZ and SlmA which are part of the divisome complex (Mahone and Goley, 2020). A particular bacterial condensate termed aggresome is formed by a diverse collection of endogenous proteins under different stress conditions. They have been associated with the bacterial fitness (Jin *et al*., 2021), protection during desiccation (Kuczyńska-Wiśnik *et al*., 2023), cell dormancy and regrowth regulation, and with drug tolerance (Pu *et al*., 2019). In particular, regarding LLPS in Archaea, although some proteins, such as the thermostable nucleoid protein Cren7 (Banerjee *et al*., 2024)), have been proposed to form liquid droplets, further studies are required.

Similarly, many viruses harness phase separation processes to accomplish nearly every step in their infection cycles—including entry, immune evasion, genome replication, new particle assembly, and particle release. As a result, LLPS is now widely recognized as a critical mechanism that facilitates the compartmentalization of viral replication within host cells. While most research has focused on the role of BMCs in replication processes (Lopez *et al*., 2021), fewer studies have examined how viral proteins disrupt host functions (Boccaccio *et al*., 2023). Furthermore, an emerging field in the role of LLPS in viruses is related with proteins adopting condensated states such as LLPS and amyloid fibrils and their roles in pathogenesis and antiviral therapeutic strategies (Gondelaud *et al*., 2023).

Most viral proteins undergoing LLPS are associated with the so-called viral factories (VF), also known as inclusion bodies, replication organelles, viral replication compartments, transcription–replication complexes, virosomes, or viroplasms (Lopez *et al*., 2021). These VF are mostly cytoplasmic spots which concentrate viral proteins and nucleic acids where viral replication and assembly is in this way optimized. Cytoplasmic location shows some exceptions such as those members of the Orthomyxoviridae and Bunyavirales order which establish replication organelles in the nucleus or in association with the Golgi complex. These VF not only serve as crucial centers for optimizing viral replication—through selective uptake or exclusion of host cellular components—but also contribute to immune evasion and escape antiviral defense pathways by trapping host proteins involved in innate immune responses. This process has been described for several viruses including Nipah virus, respiratory syncytial virus, measles virus, and SARS-CoV-2 (Li *et al*., 2022). Other viral proteins undergoing LLPS have been associated with host cell entry taking control of the host protein disaggregation pathway (Banerjee *et al*., 2014).

In addition, virus infection frequently implicates the inactivation or alteration of cellular condensates to facilitate immune escape. Among other viral components, LLPS proteins are key factors that disrupt the assembly and function of protective BMCs. An important antiviral response is the production of interferon downstream of the assembly of specific BMCs containing immunostimulatory DNA and cyclic GMP-AMP synthase (cGAS), termed cGAS foci. The formation of cGAS foci can be efficiently interrupted by viral LLPS proteins, which phase-separate and sequester activating DNA. Kaposi sarcoma-associated herpesvirus (KSHV) ORF52, also known as KSHV inhibitor of cGAS (KicGAS) and VP22 from several other viruses are emerging examples (reviewed in (Boccaccio *et al*., 2023)). The identification of novel viral LLPS will contribute to understanding viral mechanisms to escape the immune surveillance mediated by cellular BMCs.

In this work, we present a hand-curated database of proteins undergoing LLPS from bacteria, archaea, and viruses. To expand this dataset, we performed sequence similarity searches to identify closely related homologues, resulting in a comprehensive collection of over 5,000 proteins—51% from Bacteria, 1% from Archaea, 28% from Viruses, and 20% from Eukaryotes. These datasets are available to download from the web server.

### Dataset Construction

BAV-LLPS relies on three hand-curated datasets of LLPS proteins coming from bacteria, archaea and viruses. These sets contain proteins with experimental evidence of LLPS process and/or evidence of MLOs and biocondensate formation. Each entry of this curated dataset offers biological, sequence and structural information for each protein (see below). Each of these curated datasets have been extended using sequence based similarity searches using BLAST to recruit close homologues (e-values < 1.10^−5^). BLAST searches have been restricted to find homologous proteins for each taxonomic group of the curated dataset and in Eukaryotes. Moreover, we gathered together sequences coming from main available LLPS databases (PhaSepDB (You *et al*., 2020), LLPSDB (Wang *et al*., 2022), DrLLPS (Ning *et al*., 2020), PhasePro (Mészáros *et al*., 2020), and CD-Code (Rostam *et al*., 2023) with different curation levels provided from the corresponding databases. This set of curated LLPS mainly contains proteins from eukaryotic organisms (98%) and was used to further search for homologous proteins using our curated dataset coming from bacteria, archaea and viruses. As disorder content has been reported to be critical to LLPS process (Fonin *et al*., 2022; Tesei *et al*., 2021), we used Aiupred (Erdős and Dosztányi, 2024) and AF-RSA (Piovesan *et al*., 2022) to predict disordered regions, Fuzdrop (Hardenberg *et al*., 2020), ParSev2 (Wilson *et al*., 2023) and catGRANULE 2.0 (Monti *et al*., 2025) to predict LLPS per site probability and AlphaFold 2 models from AlphaFold database (https://alphafold.ebi.ac.uk/) to predict 3D models of each curated protein. When curated proteins were missing in AlphaFold database we obtain the Alpha Fold 2 model using ColabFold (Mirdita *et al*., 2022).

### Website

The web server allows users to search the entire database or to explore the three different datasets of LLPS proteins (Figure 1A). The first searchable field (“Curated database”) allows exploration of the curated dataset (94 curated proteins) deploying three tables (each for bacteria, archaea and viruses respectively). Each table contains biological information for each entry (Figure 1B). By clicking on a given protein entry, the web server opens an additional window containing sequence and structural information, along with predicted disorder and LLPS propensity per site (Figure 1C, D and E). Each of these parameters can be displayed and mapped by the user in the sequence and on the structural model using clickable buttons with tooltips that appear when hovering the mouse over the clickable buttons.

**Figure 1.**
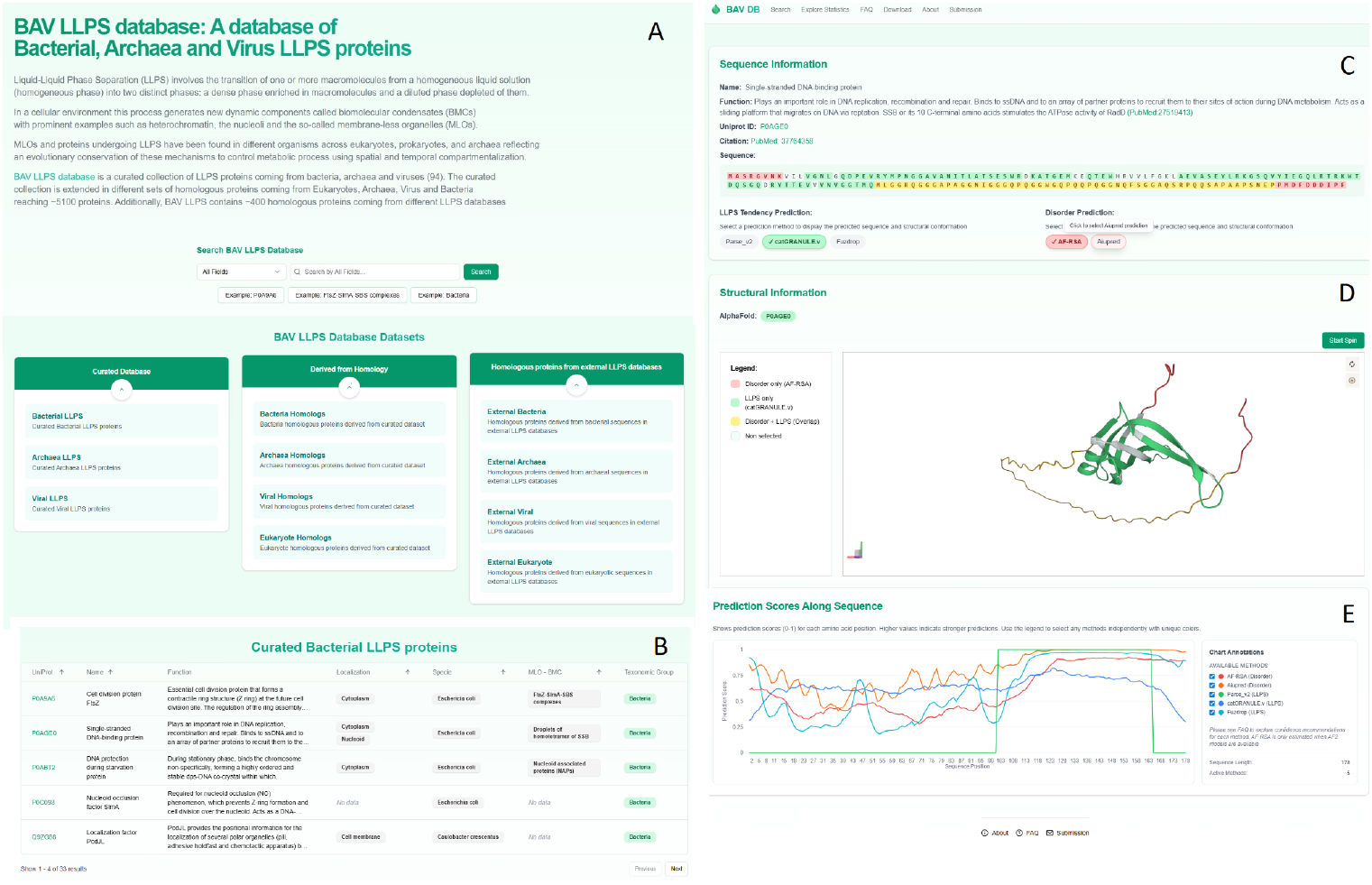
BAV-LLPS contains three main datasets: the hand-curated, homologous proteins derived from sequence similarity searches and finally a dataset of homologous proteins found in other LLPS databases (panel A). Each dataset can be viewed displaying biological information (panel B). Proteins contained in the different datasets can be accessed to explore their sequence (panel C) and structural model (panel D) along with different LLPS and disorder prediction scores that can be displayed along sequence position (panel E). Disorder and LLPS propensities per position (panel E) helps users to decide prediction confidence with the help of each method recommendations (information contained in FAQ section)

The other two searchable fields correspond to datasets of homologous proteins found in bacteria, archaea, viruses and eukaryotes (“Derived from Homology”) using the proteins of the curated dataset as inputs. As mentioned above, homologous proteins found in other LLPS databases (about 9000 unique proteins) mostly from eukaryotic organisms, can be browsed in the link called “Homologous proteins from external LLPS databases”. Similar information as displayed in Figure 1B-E is displayed for entries from these datasets.

## Conclusions

This resource aims to broaden our understanding of LLPS mechanisms beyond eukaryotic systems, promoting the study of their differential properties across a wide range of organisms. It also seeks to drive the creation of specialized predictive tools to facilitate the exploration and detailed characterization of LLPS processes in various life forms, ultimately advancing both fundamental biological research and applied biomedical sciences.

## Authors disclaimer

Views and opinions expressed are however those of the author(s) only and do not necessarily reflect those of the European Union or the European Research Executive Agency (REA). Neither the European Union nor the granting authority can be held responsible for them.

